# Data-informed discovery of hydrolytic nanozymes

**DOI:** 10.1101/2020.12.08.416305

**Authors:** Sirong Li, Zijun Zhou, Zuoxiu Tie, Bing Wang, Meng Ye, Lei Du, Ran Cui, Wei Liu, Cuihong Wan, Quanyi Liu, Sheng Zhao, Quan Wang, Yihong Zhang, Shuo Zhang, Huigang Zhang, Yan Du, Hui Wei

**Affiliations:** College of Engineering and Applied Sciences, Nanjing National Laboratory of Microstructures, Jiangsu Key Laboratory of Artificial Functional Materials, Nanjing University, Nanjing, Jiangsu 210023, China; Hubei Key Laboratory of Genetic Regulation and Integrative Biology, School of Life Sciences, Central China Normal University, No. 152 Luoyu Road, Wuhan, Hubei 430079, China; Jiangsu Key Laboratory of Druggability of Biopharmaceuticals, State Key Laboratory of Natural Medicines, School of Life Science and Technology, China Pharmaceutical University, Nanjing, Jiangsu 211198, China; College of Chemistry and Molecular Sciences, Wuhan University, Wuhan, Hubei 430072, China; State Key Laboratory of Electroanalytical Chemistry, Changchun Institute of Applied Chemistry, Chinese Academy of Sciences, Changchun, Jilin 130022, China; University of Science and Technology of China, Hefei, Anhui, Hefei 230026, China; Collaborative Innovation Center of Advanced Microstructures and Institute of Materials Engineering Nanjing University, Nanjing, Jiangsu 210093, China; State Key Laboratory of Analytical Chemistry for Life Science, School of Chemistry and Chemical Engineering, Nanjing University, Nanjing, Jiangsu 210023, China; Chemistry and Biomedicine Innovation Center (ChemBIC), Nanjing University, Nanjing, Jiangsu 210023, China

**Keywords:** data-informed, rational design, metal–organic framework, hydrolase mimic, nanozymes

## Abstract

Nanozyme is a collection of nanomaterials with enzyme-like activity but exhibits higher environmental tolerance and long-term stability than their natural counterparts. Improving the catalytic activity and expanding the category of nanozymes are prerequisites to complement or even supersede natural enzymes. Specifically, a powerful hydrolytic nanozyme is demanded to degrade the unsustainable substance which natural enzymes hardly achieve. However, the development of hydrolytic nanozymes is still hindered by diverse hydrolytic substrates and following complicated mechanisms. Here, we apply two strategies which are informed by data to screen and predict catalytic active sites of MOF (metal–organic framework) based hydrolytic nanozymes. One is to increase the intrinsic activity by finely tuned Lewis acidity of the metal clusters. The other is to adjust the volume density of the active sites by shortening the length of ligands. Finally, we construct a Ce-FMA-MOF-based hydrolytic nanozyme with robust cleavage ability towards phosphate bonds, amide bonds, glycosidic bonds whose energy ascend in order; and even their mixture, biofilms. This work provides a rational methodology to design hydrolytic nanozyme, enriches the diversity of nanozymes, and potentially sheds a light on the evolution of enzyme engineering in the future.

## Introduction

Exploring enzyme mimics in artificially fabricated systems is a promising strategy to overcome the instability and high cost of enzymes^1^. It can also enable us to better understand the living world^2^. However, the development of this strategy is hindered by limited knowledge on miscellaneous mechanisms and finite chemical methodology to mimic. Over the last two decades, the emergence of nanotechnology has expanded artificial enzymes into nanomaterials, which are now collectively termed as nanozymes^3–7^. Nanozymes integrate multivalent catalytic sites while retain the multifunctional repertoires of nanomaterials, such as magnetic property of Fe_3_O_4_. Thereby diverse feats in bioanalysis^4,8,9^, medical imaging^10^, therapeutics^11^, and tissue engineering^12^ have been achieved, enriching biomimetic nanozymes to a larger context.

In our recent review on nanozymes^3^, we summarized an exponential growth in the number of publications on nanozymes, demonstrating the fast expansion of this field. However, the breadth of enzymatic reactions that has been explored is, to date, rather limited. Specifically, a detailed analysis of these papers showed that only a small fraction (7.1%) focused on hydrolytic enzyme mimics, while the majority (92.9%) focused on redox enzyme mimics (Figure 1a). To fully exploit nanozymes, it is demanded to widen the category that nanozymes can mimic and study them in-depth to compensate this 13-fold difference in the number of studies on hydrolase and redox enzyme mimics.

**Figure 1.**
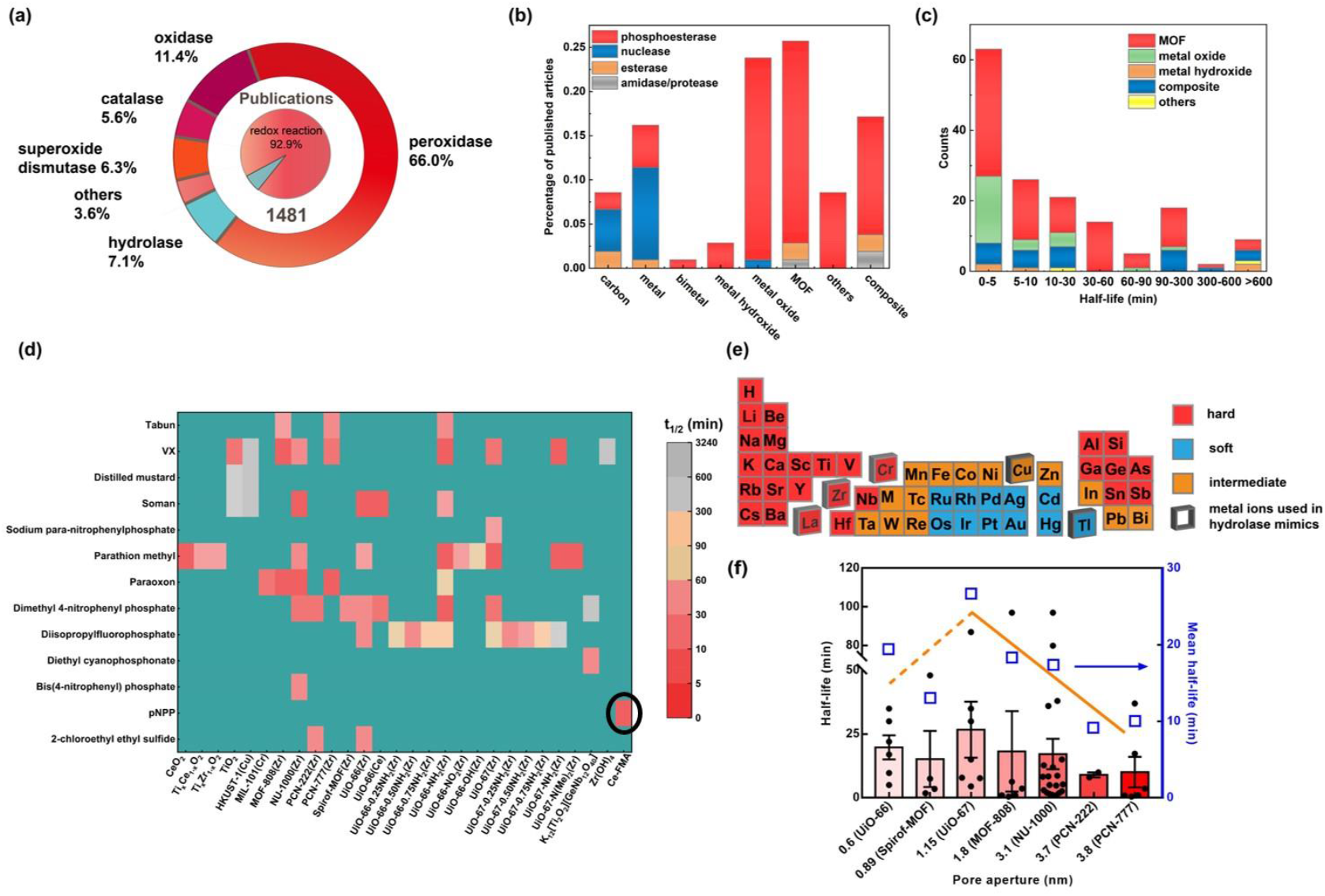
The overall statistical analysis of hydrolytic nanozymes. (a) Proportions of different types of nanozymes. Raw data were adapted from Reference 3. (b) Stacked histogram of the publication frequency distribution of studies on different types of material related to each type of hydrolase. (c) Stacked histogram of the number distribution of each material based on its hydrolytic half-life. (d) Heat map of the half-lives of various phosphatase-like nanozymes and their corresponding substrates. Each half-life is presented as the mean unless there is only one sample. The circled one represents the data-informed Ce-FMA in this work. (e) Periodic table of metal elements mapped by hard-soft-acid-base theory. The red indicates hard Lewis acid, the blue indicates soft Lewis acid, and the yellow indicates the intermediate Lewis acid. Titled squares represent metal compositions that have been reported with hydrolase-like activity. (f) Half-lives of reported Zr-based MOFs versus their pore aperture; the mean values are shown as blue empty squares. The number associated with each MOF on the x-axis represents the pore/window aperture^14, 29–34^. Each half-life is presented as the mean ± standard deviation.

Biologically, hydrolases are involved in ubiquitous bio-transformations with substrates such as esters (esterases), phosphoesters (phosphoesterases), amides (amidases and proteases), and carbohydrates (glycosidases); see Table S1. The broad substrate scope of hydrolases enables them to function in various biological processes, such as nerve impulse transmission (*e.g.*, acetylcholinesterase), blood sugar balance (*e.g.*, glycogen phosphorylase), dissimilation (*e.g.*, digestive enzyme), and energy transfer (*e.g.*, ATP hydrolase). Additionally, organophosphate-like nanozymes can degrade organophosphorus pesticides/chemical warfare agents (such as soman, sarin, and tabun) by breaking the P-X bond (X=O, F, CN, *etc.*)^13^, hopeful to be embedded into protective masks. Despite the encouraging achievements, to date, only a few such nanozymes have been developed, which can be attributed, at least in part to that the current design strategies are heavily relying on integrating natural active site moieties (such as Zn^2+^-coordinated complexes) within/onto nanomaterials^5, 14–19^. The specificity and regulation of natural active sites restrict the further evolution of these nanozymes in abiotic environment or specific microenvironment, which may lead to non-optimal reactivity and thus an unsuitable replication template for new-to-nature catalysis^20–22^. Moreover, the diversity of substrates and relatively limited knowledge on the catalytic mechanisms of hydrolases give additional obstacles towards the design of new hydrolytic nanozymes. To overcome these challenges and expand nanozyme functionality beyond nature’s repertoire, herein, we successfully identified two key factors to design MOF (metal–organic framework) based hydrolytic nanozymes through a data-informed analysis of published hydrolase-like nanozymes based on our recent comprehensive review^3^.

Data-informed analysis is an approach which balances between expertise and understanding of information, thus draws key information and then derives insights from structured and unstructured data^23^. In this work, 105 research papers describing hydrolytic nanozymes is screened from 1481 research papers we reviewed (the list of papers we covered is supplemented as an endnote file). The breadth and the efficiency heat map of hydrolytic nanozymes (Figure 1d) indicated MOF as a good scaffold to incorporate active sites. Further, two factors were informed through data of which the metal ions were suggested as hard Lewis acid (*i.e.*, Ce^4+^) for higher affinity with substrates while the ligand was deduced as fumaric acid in that a shorter ligand could increase the volume density of the active sites. The optimized hydrolytic MOF nanozyme was prepared experimentally and shown to possess excellent phosphatase-, protease-, and glycosidase-like activities among current nanozymes. Moreover, the optimized hydrolytic nanozyme even successfully degraded biofilms under mild conditions. This work provides a general methodology to design hydrolytic nanozymes. And the abiotic active sites we derived may conversely shed a light on enzyme engineering in the future.

### Data-informed strategy to identify hydrolase-like candidate materials

The course to develop a potent hydrolytic nanozyme is: 1) selecting a suitable Scaffold to embed/design highly active sites; 2) identifying a highly reactive site to activate hydrolytic substrates; 3) performing experiments to fabricate designed materials. We envisioned the suitable Scaffold to embed active sites for nanozymes can be deduced from data classified by varying material category. Therefore, we first plotted the publication frequency distribution of studies on different types of material related to each type of hydrolase in Figure 1b. Four types of hydrolase, namely, phosphoesterase, nuclease, esterase, and amidase/protease (there were no glycosidase mimics when this work was initiated in mid-2018) were sorted which were mimicked by carbon, metal, bimetal, metal hydroxide, metal oxide, MOF, composite, and others. Notably, reports on MOFs, which are crystalline materials consisting of metal clusters coordinated by organic ligands^24–26^, and metal oxides were the most numerous, indicating these two as scaffold candidates to mimic hydrolases. On the other hand, half-life (t*1/2*, the time needed to achieve 50% conversion) which was presented as the maximum kinetic parameter among collected data, is grouped in Figure 1c to determine which scaffold is more likely to achieve faster catalytic speed. MOFs quickly draw a specific focus because of their much higher representation in the literature than other materials with a sub-10 min half-life. Consequently, we inferred that MOFs are optimal candidates to mimic hydrolases.

Having confirming MOFs as a potent scaffold, our next goal was to deduce highly active sites of MOFs. A kinetic heat map that displayed the half-lives of various phosphorylated substrates treated with nanozymes was shown in Figure 1d. Intriguingly, though dozens of materials were reported to mimic hydrolase, the metal components were within five elements, namely zirconium (IV), cerium (IV), chromium (III), copper (II), and titanium (IV), see the tilted box in Figure 1e. Moreover, nanozymes consisted of metal ions with hard Lewis acidy tend to have faster half-lives, such as zirconium (IV), the major (17/20) component of the hydrolytic MOFs, manifesting hard Lewis acid as effective active sites. Mechanistically, a hard Lewis acid, such as a high-valence metal ion, can easily activate a carbonyl or phosphoryl group by accepting an electron lone pair from the oxygen and drawing electron density away from the double bond, leading to a greater positive charge on—and thus increasing the electrophilicity and reactivity of—the central carbon or phosphorus^28,29^. Thus, Lewis acid metal ions are preferred active sites in a MOF scaffold for mimicking hydrolases, based on both an analysis of previously published data and hard-soft-acid-base theory.

Another impact factor in MOF scaffolds is the ligand which controls over the metal spacing/volume density of active sites and moreover affects the final pore/window aperture of MOFs. Since different connectivity between the same metal ion and ligand can yield MOFs with different topology structures and pores/windows, we classified the MOFs in the abovementioned papers by their pore/window aperture instead of ligand and plotted their kinetic data (half-lives) in Figure 1f. For MOFs with pore widths above approximately 1 nm, larger pores lead to faster reaction rates (*i.e.*, shorter half-lives) due to easier substrate diffusion into the catalytic interior of the MOF (solid yellow trend line, Figure 1f)^27^. Below pore widths of approximately 1 nm, however, this trend is reversed. We attributed this reversed trend to the increased volume density of active site as shorter ligand means greater vicinity of metal clusters^28^. Of the MOFs in which this trend is observed, UiO-66 and UiO-67 are of particular interest, as these two isostructural MOFs differ only in the length of their structural ligands (shorter ligand benzene-1,4-dicarboxylic acid (BDC) in UiO-66 and longer ligand biphenyl-4,4’-dicarboxylate (BPDC) in UiO-67; Figure S1). In this regard, we selected fumaric acid (FMA, Figure S2) as a candidate ligand because its length is shorter than BDC but able to form/construct a homologue of UiO-66. Of note, there currently are no available data showing that an FMA contained MOF possesses phosphoesterase-like activity.

Based on the above analyses, we selected high-valence, strong Lewis acid ions (Zr^4+^, Ce^4+^, and Hf^4+^, the three tetravalent metal ions with the strongest Lewis acidity in Table S2) and FMA to construct a homologue MOF of UiO-66.

### Material synthesis, optimization, and characterization

#### Synthesis of Zr-FMA, Hf-FMA and Ce-FMA

With this data analysis in mind, we fabricated several MOFs with high-valence metal ions (*i.e.*, Zr^4+^, Ce^4+^, and Hf^4+^) bridged by FMA. The aqueous synthesis of Zr-FMA and Hf-FMA was performed according to previous work^35^, while the aqueous synthesis of Ce-FMA was conducted in batches of trials by varying the temperature, reaction time, and modulators since so far no aqueous synthesis of Ce-FMA was reported. First, Ce-FMA was synthesized without modulators to exam which temperature and reaction time could yield crystalline products. Products were collected after 10 min of stirring at room temperature and subsequent stirring for a longer time under reflux (around 105 °C). X-ray diffraction (XRD) results confirmed that the crystal structure remained uniform from 10 min at room temperature to 180 min under reflux (Figure S4a); additionally, the activity of these MOFs towards the hydrolysis of *para*-nitrophenyl phosphate (pNPP) remained almost identical over reaction time (Figure S4b). Therefore, stirring at room temperature for 10 min without the additional high-temperature step was applied in subsequent trials to evaluate the effect of modulators. An overall survey of modulators on hydrolase-like MOFs from the 105 published papers is summarized in Table S4, denoting that the yield^30^, degree of crystallinity^30^, morphology/size^17^, and presence of defects^36^ of the synthesized MOFs could be regulated by the modulators^37^, ultimately resulting in differences in the catalytic performance. We selected acetic acid (AA), formic acid (FA), and trifluoroacetate (TFA) (see their structures in Figure S3 and the synthetic details in Table S5) for further study because they possess similar fatty structures to that of FMA. The XRD results, transmission electron microscopy (TEM) images, and the Brunner-Emmet-Teller (BET) surface area of Ce-FMA obtained under each condition are displayed in Figures S5, S6, S7 and Table S6. For optimization, modulator varied Ce-FMA MOFs were evaluated based on the catalytic performance in phosphatase-like activity assays with the phosphatase substrates pNPP and bis-*p*-nitrophenyl phosphate (BNPP), as discussed below.

#### Optimization of the metal ions and ligands

In addition to Zr-FMA, Hf-FMA, and Ce-FMA, we fabricated Zr-BDC, Hf-BDC, and Ce-BDC to investigate which metal ions dominate the hydrolytic activity and whether increasing the volume density of active sites by shortening the length of ligand increase the activity. Having confirmed the identical phase structure of six MOFs by XRD, as shown in Figure S8a, we then tested phosphatase-like activity towards pNPP. Neither Hf-BDC nor Hf-FMA showed prominent phosphatase-like activity, as shown in Figure S8b and S8c, and these samples therefore served as null examples of Hf-based hydrolytic nanozymes. We then compared Zr- and Ce-based MOFs linked by FMA and BDC. Consistently, the Ce-based systems demonstrated greater catalytic activity than the Zr-based systems (Figure S8d and S8e), and nonactivated Ce-FMA demonstrated higher activity than activated Ce-BDC (Figure S8f), even though the BET surface area of Ce-FMA was approximately 4 times smaller than that of activated Ce-BDC (120.4338 m^2^/g versus 516.9987 m^2^/g, respectively; Table S6). This difference between systems with different metal ions and ligands, for one thing, demonstrates our hypothesis that a shorter ligand is beneficial to yield higher volume density to the active sites; for another, indicates that the increased Lewis acidy of Ce makes the 4f orbital well suited to hybridize with the P-O bond and thus better stabilizes the pentavalent phosphate intermediate for nucleophilic attack. Both factors collectively make the Ce-FMA as particularly well-suited for a hydrolase-like catalyst^38^.

#### Characterization of an optimal Ce-FMA

An overall comparison of the various synthetic conditions for Ce-FMA was conducted via phosphatase-like activity assays (described in supplementary information) with the substrates pNPP and BNPP. An FA-to-FMA molar ratio of 20 was determined to be the best synthetic condition according to the phosphatase-like activity (discussed in the following catalytic performance section), and this optimized structure was referred to as Ce-FMA-FA-20-RT. As illustrated in Figure 2a, the [Ce_6_O_4_(OH)_4_]^12+^ clusters are arranged as cubic close packing and linked by twelve FMA^2−^, yielding the formula as [Ce_6_O_4_(OH)_4_(FMA)_6_] in Ce-FMA. The XRD patterns in Figure 2b confirmed consistent crystal structure with simulated results^40^. TEM images revealed that this modulated MOF was approximately 180-210 nm in diameter with slight aggregation (Figure 2c). X-ray photoemission spectroscopy (XPS) deconvolution of the peaks of Ce 3d in Figure 2d indicates a proportion of 82.7% Ce^4+^ and 17.3% of Ce^3+^, making the final valence of 3.8. A typical three-stage thermal behaviour determined by thermogravimetric analysis was also exhibited (Figure 2e), suggesting a similar thermal behaviour with Ce-BDC^39^. Fourier transform infrared spectroscopy (FTIR) in Figure 2f also demonstrates the remained trans structured C=C in MOFs derived from FMA.

**Figure 2.**
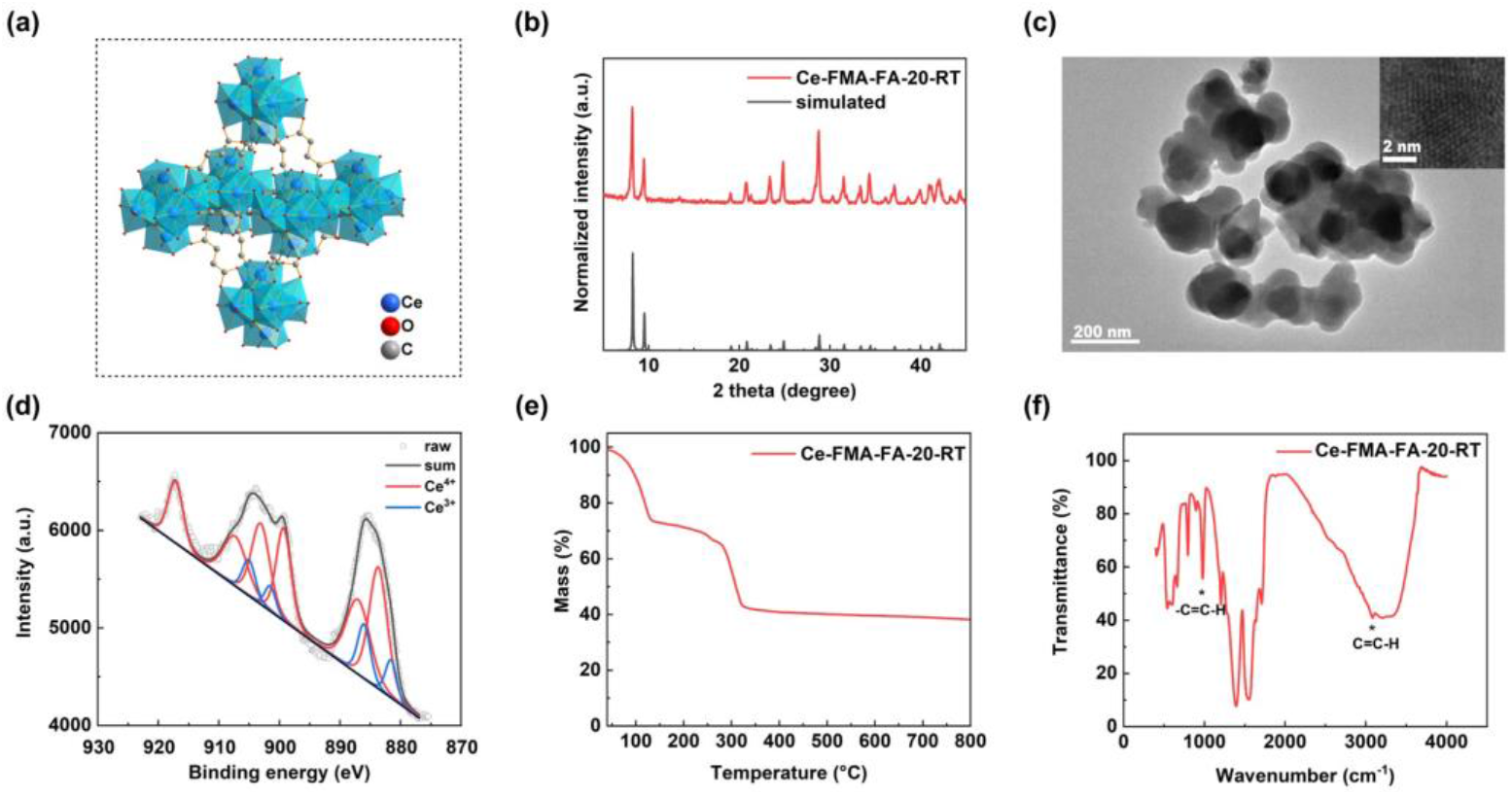
Characterization of Ce-FMA modulated by FA with an FA-to-FMA molar ratio of 20. (a) Structure of Ce-FMA. Ce blue, O red, C gray; hydrogen atoms are omitted for clarity. (b) XRD pattern of Ce-FMA-FA-20-RT (the black lines at the bottom are simulated pattern from reference 39). (c) TEM images of Ce-FMA-FA-20-RT at different magnifications. (d) XPS spectrum of the Ce 3d signals of the Ce-FMA-FA-20-RT. The red curves correspond to Ce^4+^ which are deconvoluted into three Voigt doublets and the blue curves correspond to Ce^3+^ which are deconvoluted into two Voigt doublets. The black curve is summed by every deconvoluted peak. (e) TG curve of Ce-FMA-FA-20-RT analysed under air flow. (f) FTIR spectrum of Ce-FMA-FA-20-RT.

### Catalytic performance

Different substrates (listed in Table S3) towards four main hydrolases were applied to evaluate the catalytic performance of Ce-FMA. Since the phosphoester bond is the most active bond among hydrolytic bonds, we studied phosphatase-like activity first and used the activity data to verify the optimal synthetic conditions (confirming the modulator and the dosage) of Ce-FMA. Then we explored Ce-FMA to hydrolyse planar amide bond in bovine serum albumin (BSA) which is more hierarchal in structures and difficult to cleave. With these success, we next investigated glycoside bonds cleavage effect in chitosan. In summary, except for carbonate esterase, Ce-FMA is able to mimic the other three types of hydrolase.

#### Phosphatase-like activity

Despite the variation in all the phosphorylase substrates previously reported (summarized in Figure 1d), the catalytic mechanism was consistent with Lewis acid-activated cleavage of the P-X bond (X=O, F, CN, *etc.*). For phosphorylated substrates, as illustrated in Figure 3a, the reaction starts with the nucleophilic addition of an undercoordinated M-OH (M refers to metal) after the substrate was activated by metal cluster in MOF, forming a pentacoordinated phosphorus intermediate. And then, the intermediate decomposed via the elimination of alcohol^40^. Since most of the organophosphorus are neurotoxic, for analytical efficiency and safety, we selected pNPP and BNPP as phosphatase substrates instead of the other organophosphorus compounds. As shown in Figure 3a, both pNPP and BNPP produced yellowish 4-nitrophenol after phosphoester bond cleavage, allowing us to measure the reaction rate quantitatively by recording the absorbance at 400-405 nm. Usually, pNPP is used as a substrate of alkaline phosphatase (ALP), an enzyme that can convert phosphate compounds (*i.e.*, pyrophosphate) to free phosphate and thus accelerate mineralization in hard tissue formation. Likewise, both Ce-FMA and ALP exhibited pH-dependent activity and achieved their maximum activities at pH 10.0 (Figure S10). Both the mass activity and surface area-normalized activity towards pNPP and BNPP were taken into consideration to optimize the synthetic conditions of Ce-FMA (Figure 3b–3e) since it is convenient to use mass activity to evaluate cost, but more consistent with the active sites by surface area-normalized activity. In general, Ce-FMA synthesized with an FA-to-FMA molar ratio of 20 resulted in the highest phosphatase-like activity.

**Figure 3.**
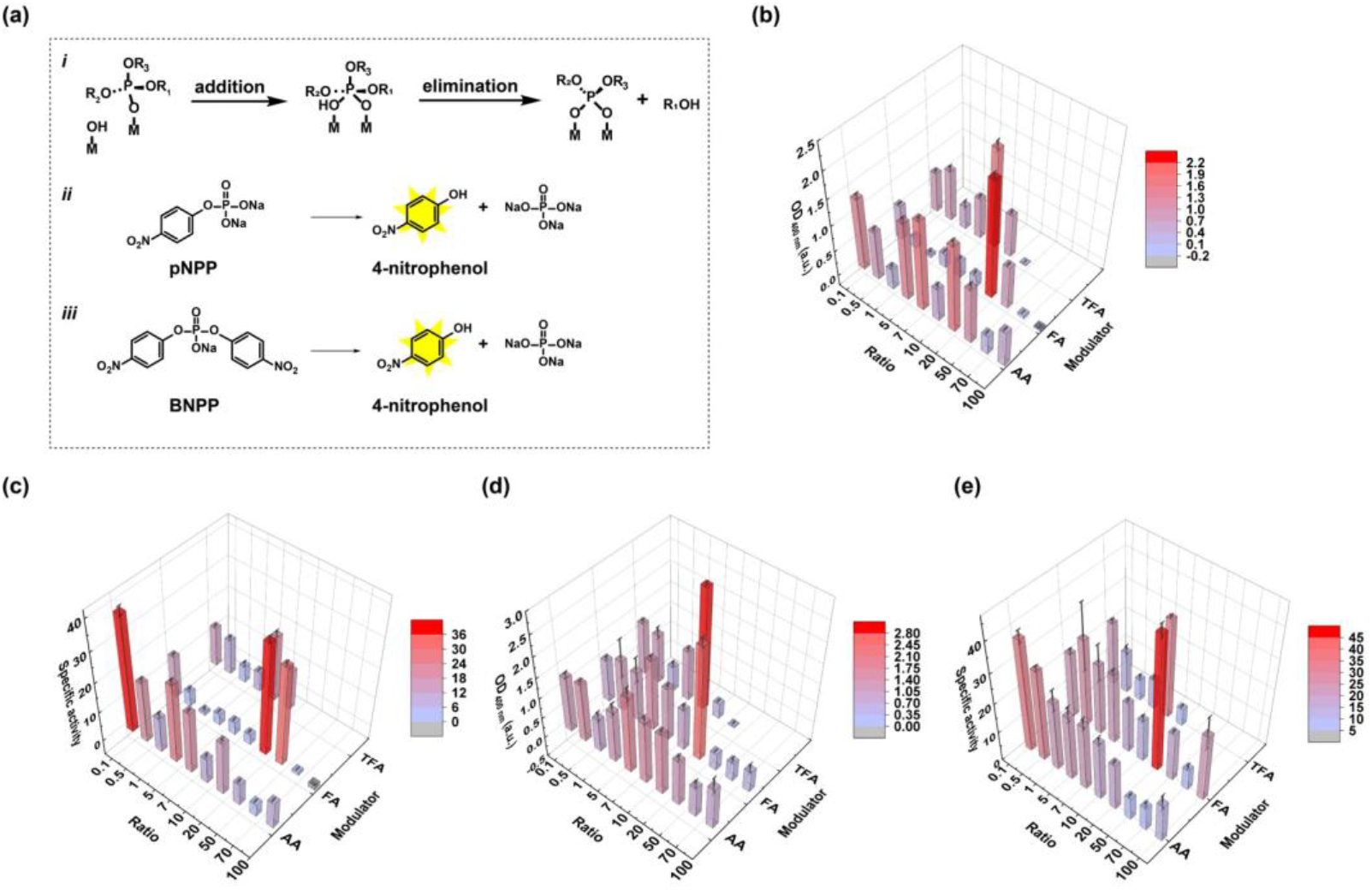
Phosphatase-like activity of Ce-FMA. (a) Schematic of the hydrolytic mechanism and the detection principles of pNPP and BNPP. Both pNPP and BNPP produce yellowish 4-nitrophenol after breaking phosphomonoester and phosphodiester bonds. (b) and (c) 3D histogram maps of the mass and specific ALP-like activity of Ce-FMA with the substrate pNPP. (d) and (e) 3D histogram maps of the mass and specific phosphodiesterase-like activity of Ce-FMA with the substrate BNPP. The activity is indicated by the colour gradient. Data are processed by removing the self-hydrolysis effect and the initial background of nanozymes. Data are presented as the mean ± standard error after removing the maximum and minimum, n=8.

After successfully demonstrating the phosphomonoesterase-like activity of Ce-FMA, we next investigated the phosphodiesterase-like activity of Ce-FMA by using BNPP as the substrate. However, while the activity for BNPP hydrolysis increased with the pH, similar to that for pNPP hydrolysis, the optimized pH was 9.0, as shown in Figure S11a. This may be because the rate-limiting step becomes substrate binding when pH is within 9.0 to 10.0 rather than nucleophilic attack when pH is within 7.0 to 9.0^41^. On the other hand, BNPP underwent a fast spontaneous hydrolysis at alkaline solutions or high concentration (Figure S11b and S11c), indicating easier cleavable property than pNPP. Therefore, Ce-FMA with good activity towards pNPP (modulated by FA with an FA-to-FMA ratio of 20) was still active when it hydrolyzed BNPP (Figure 3d and 3e) and even showed faster kinetics.

In addition to pNPP and BNPP, we also tested whether optimized Ce-FMA-FA-20-RT could cleave the phosphate backbone in DNA as BNPP is usually considered the simplest model substrate of DNA. However, no fragments of DNA were detected (Figure S12). This difference could be rationalized by the two excellent leaving groups in BNPP which is not present in DNA.

In summary, the half-life of pNPP hydrolysis under the action of Ce-FMA-FA-20-RT (within 2 min; Figure S8f) is marked in the heat map in Figure 1d to have a comprehensive overview. Remarkably, we did not apply the normally required co-catalyst such as polyethylenimine^38^ or *N*-ethylmorpholine^27^ for this data-informed Ce-FMA MOF.

#### Protease-like activity

Given the breadth of biologically and chemically relevant hydrolase reactions and mechanisms, we were curious whether Ce-FMA could mimic other hydrolytic enzymes beyond phosphoesterases. Thus, we next evaluated the catalytic performance of Ce-FMA as a protease. Hydrolysis of amide/peptide bonds in proteins is more challenging than that of phosphate bonds because the phosphate linkage is more easily cleaved. In contrast to the autolysis of phosphate bond (such as phosphodiester linkage in BNPP, as shown in Figure S11b and S11c), the autolysis of amide/peptide bonds has a half-life of 350 years under physiological pH and temperature^42^.

We applied bovine serum albumin (BSA) as a substrate to test the protease-like activity. BSA is a commonly used and stable globular protein with 585 amino acids (66.4 kDa molecular weight). To monitor the hydrolysis of BSA, gel permeation chromatography (GPC) with UV detection at 280 nm was used. The peak corresponding to BSA at 34-36 min decreased gradually as the reaction proceeded in PBS (pH 7.2-7.4) at both 60 °C and 37 °C, see the lower curves in Figure 4a and 4b; while there is no noticeable self-degradation product in the absence of Ce-FMA-FA-20-RT, see the upper curves in Figure 4a and 4b. A 100% conversion rate was achieved after 36 h and 7 d respectively at 60 °C and 37 °C in Figure 4d and 4e. We then compared the hydrolysis efficiency of our MOF with trypsin (comparison was conducted at 37 °C) and MOF-808 (comparison was conducted at 60 °C) ([Zr_6_O_4_(OH_4_(BTC)_2_-(HCOO)_6_] (BTC: benzene-1,3,5-tricarboxylate) because trypsin is a typical protease while MOF-808 was reported as both good protease-^30^ and phosphatase-like nanozyme^43^. For trypsin, a conversion rate of 100% was achieved after 24 h as shown in Figure S13, demonstrating 7 times higher efficiency than Ce-FMA-FA-20-RT (37 °C, 7 days) but ten thousand more costly than nanozyme. For MOF-808, it achieved no more than 50% conversion after 24 h, while the conversion of Ce-FMA-FA-20-RT reached 75.54% (Figures 4d, S14c, and S14d), even though the surface area of MOF-808 is more than 8 times greater than that of Ce-FMA-FA-20-RT (1017.8893 m^2^/g versus 120.4338 m^2^/g, respectively, Table S6), further confirming the advantage of Ce^4+^ over Zr^4+^ in the active site of hydrolytic MOFs.

**Figure 4.**
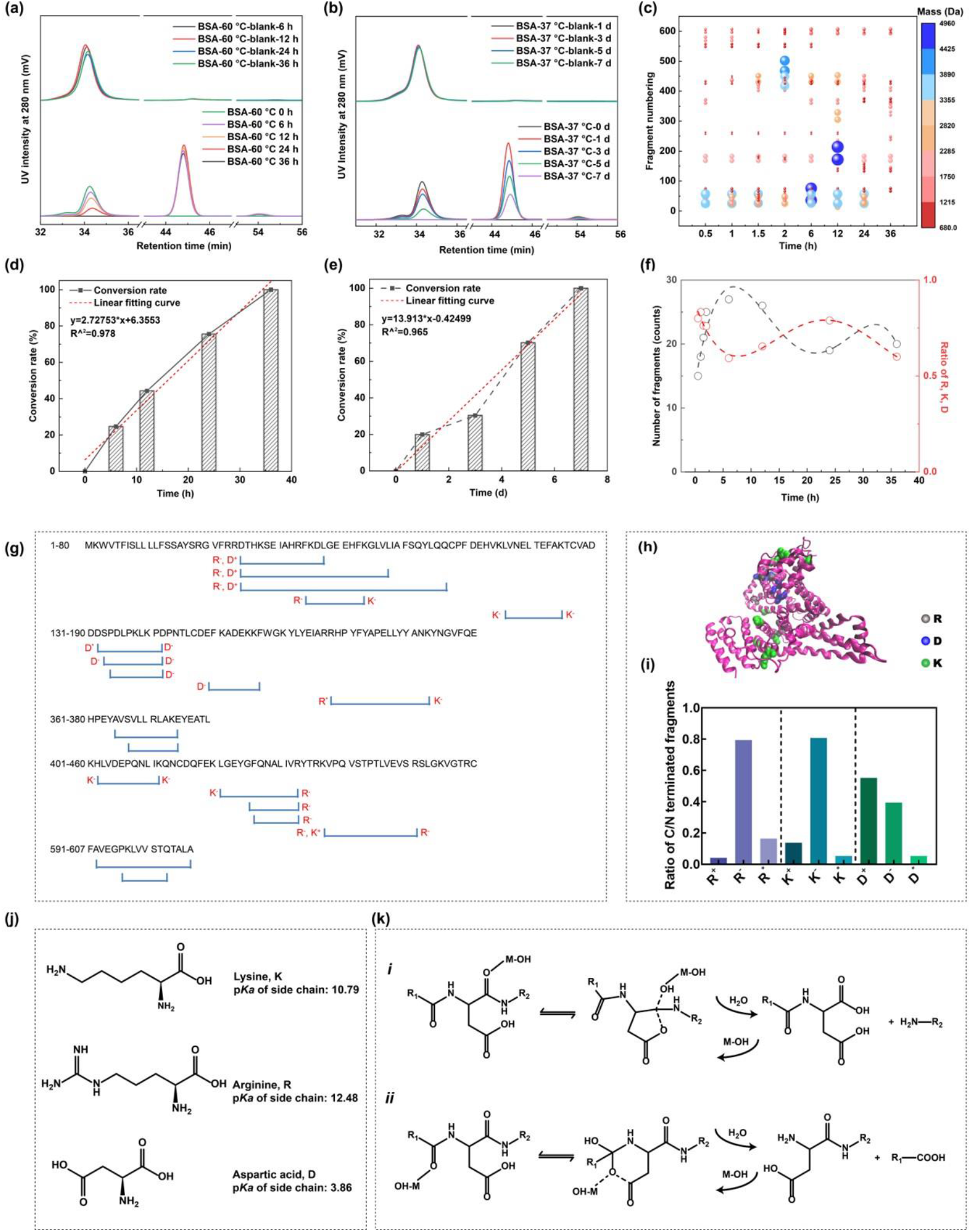
Protease-like activity of Ce-FMA-FA-20-RT. (a) GPC profiles of BSA with/without Ce-FMA-FA-20-RT in PBS (pH 7.4) at 60 °C. (b) GPC profiles of BSA with/without Ce-FMA-FA-20-RT in PBS (pH 7.4) at 37 °C. (c) Fragment numbering and mass of fragments of Ce-FMA-FA-20-RT degraded BSA collected at different time. The mass is indicated by the colour gradient and the size of the bubble. (d) and (e) Time dependent conversion rate of BSA corresponding to (a) and (b), which is calculated by the peak area integral of BSA. (f) Number of fragments and ratio of fragments with indicated termini versus reaction time. (g) Sequence coverage of BSA and Ce-FMA-FA-20-RT cleaved peptides after 24 h. (h) Visualization of BSA and the surface distribution of cleaved R, K and D after 24 h at 60 °C (PDB code: 6QS9). (i) Histogram of the frequency distribution of R, K and D. Specifically, each cleavage site is classified as N-terminus (R^+^, K^+^ and D^+^), C-terminus (R^−^, K^−^ and D^−^) or both (R^*^, K^*^ and D^*^). (j) Chemical structures of K, R and D and the p*Ka* values of their side chains. (k) Mechanisms of cleavage at D^−^ (*i*) and D^+^ (*ii*) termini, showing both of them are thermally stable to form the cyclic anhydride and imide intermediates, adapted from reference 44.

However, peaks ascribed to degradation products (44-46 min and 53-55 min) were not consistent with the peak of tryptophan (referred as possible final hydrolysis products) whose peak appeared at 93 min in Figure S15, indicating products with molecular weight larger than tryptophan formed. To obtain the exact molecular weight of the hydrolysed fragments and to identify possible cleavage sites, we further carried out electrospray ionization mass spectrometry (ESI-MS) analysis. As shown in Figure 4c, during the process of hydrolysis, the molecular weight of cleaved fragments ranged from 699 to 4764 Da, composed of 6 to 41 amino acids. Notably, there were no fragments larger than 5000 D*a* even though we collected the sample at the initial process of hydrolysis (30 min), indicating only small fragments can desorb from Ce-FMA-FA-20-RT. Moreover, the fragments were finally digested into fragments with 6 to 12 amino acids at 36 h, which is alike length with natural protease^45^. Coverage of BSA sequence by Ce-FMA-FA-20-RT cleaved peptides from 0.5 h to 36 h indicates arginine (R), lysine (K), and aspartic acid (D) are the three main cleavage sites, see Figures 4f, 4g and S16. The ratio of selectivity (number of fragments with R or K or D/number of total fragments) are plotted in Figure 4f. Reasonably, the ratio of selectivity decreases when the number of fragments increases but increases if the number of fragments decrease. Moreover, the cleaved sites, plotted in Figure 4h, are located on the outer surface of BSA, suggesting a crucial interaction between the outer surface of BSA and Ce-FMA-FA-20-RT which can also be proved by TEM images in Figure S9e and S9f.

The high protease-like activity of Ce-FMA-FA-20-RT derives from Lewis acid activation mechanism. Similar to the activation of organophosphorus, the Lewis acid (*i.e.*, Ce^4+^) activates carbon by polarizing the peptide bond after coordinating with the amide oxygen, thus enhancing the affinity for nucleophile attack to break the amide bond. We ascribed the selective cleavage on sites of R, K and D to three reasons. First, the alkaline/acid groups in side chains accelerate the process of hydrolysis, see in Figure 4j. Second, the NH_2_ groups in the side chain of R and K act as H-bond donor and later form oxyanion-hole to stabilize the intermediate^46^. Last, hydrolysis of D may undergo two pathways with cyclic anhydride or imide intermediate (Figure 4k), thereby leading to an even distribution of N-terminal and C-terminal in D sites in Figure 4i. Understanding the specific interaction is helpful for future design of site-selective protease-like nanozyme and accordingly to acquire potential bioactive peptides. Moreover, since Ce-FMA-FA-20-RT showed different cleavage positions from that by trypsin (R, K, and D for Ce-FMA-FA-20-RT *vs.* R and K for trypsin), we further summarized a comparison between Ce-FMA-FA-20-RT and trypsin towards cost, catalytic efficiency and storage in Table S7. In conclusion, even though trypsin showed one order of magnitude higher catalytic efficiency than Ce-FMA-FA-20-RT, this data-informed nanozyme decreased four order of magnitudes in cost, making itself an optional alternative.

#### Glycosidase-like activity

The successful demonstration of the ability of Ce-FMA-FA-20-RT to cleave both phosphate bonds and peptide bonds encouraged us to further explore the hydrolytic activity of this material towards glycosidic bonds. Glycosidase-like nanozymes have shown great promise in nanozyme-enabled prodrug therapy in the form of glucuronide prodrugs to treat inflammation and bacterial infection^47^. To evaluate the broad substrate scope of Ce-FMA-FA-20-RT, we started with chromogenic substrates of 2-nitrophenyl β-D-galactopyranoside and 4-nitrophenyl N-acetyl-β-D-glucosaminide. As shown in Figure 5a and 5b, both of the substrates are linked by a β-glycosidic bond without/with an acetyl amino group. To our surprise, Ce-FMA-FA-20-RT is more efficient to cleave 4-nitrophenyl N-acetyl-β-D-glucosaminide than 2-nitrophenyl β-D-galactopyranoside (see in Figure 5c, 5d and Figure S17, in which a higher conversion was observed in 4-nitrophenyl N-acetyl-β-D-glucosaminide). We further applied an ion chromatography to investigate whether Ce-FMA-FA-20-RT could cleave α-1,4 glycosidic bonds in maltose and β-1,4 glycosidic bonds in lactose when neither of the two substrates have an acetyl amino group. However, there were no cleaved monosaccharide products (Figure S18). These paradoxical results drove us to investigate the effect of acetyl amino group. Thus, carboxymethyl chitosan with acetyl amino functionalized β-1,4 glycosidic bonds were investigated in alkaline environments. GPC with RID detector was used to measure the product of carboxymethyl chitosan. GPC results demonstrate that Ce-FMA-FA-20-RT is able to cleave carboxymethyl chitosan in alkaline (pH 8) environments both at 37 °C and 60 °C, see Figure 5e. More, the catalytic efficiency increased with the temperature increased as the product peak in 60 °C shifted more right than that in 37 °C (retention time 42-44 min *vs*. 40-42 min). We ascribed this to the formation of H-bonding between N of acetyl amino and Ce cluster. Additionally, the TEM morphology of Ce-FMA-FA-20-RT before and after catalysis was similar (Figure S9a-S9e).

**Figure 5.**
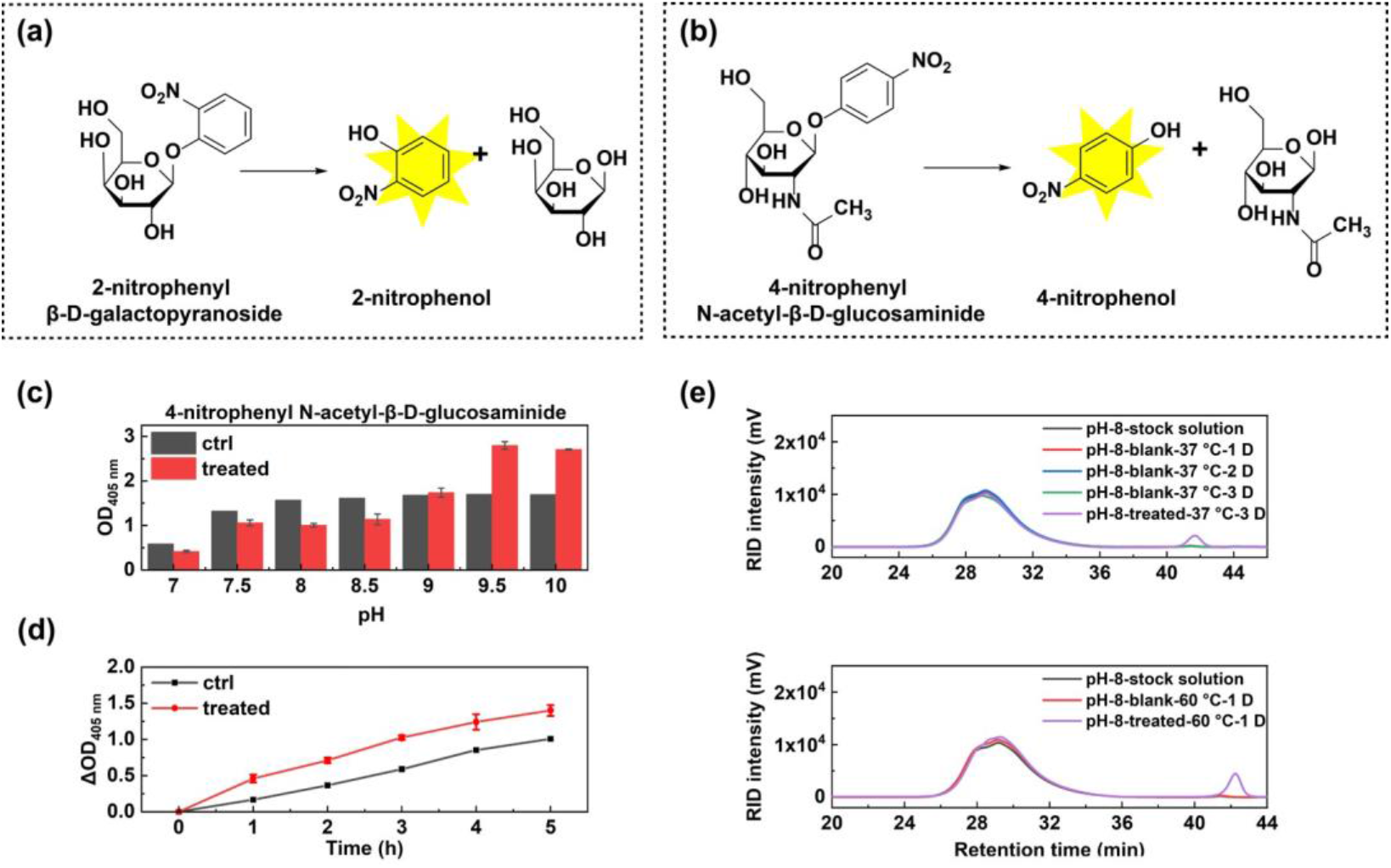
Glycosidase-like activity of Ce-FMA-FA-20-RT. (a) and (b) Schematic illustration of hydrolysis and the detection principle of 2-nitrophenyl β-D-galactopyranoside in (a) and 4-nitrophenyl N-acetyl-β-D-glucosaminide in (b). (c) The optical density at 405 nm of hydrolysed product of 4-nitrophenyl N-acetyl-β-D-glucosaminide under different pHs at 60 °C for 8 h. Data are presented as mean ± standard error of the mean (n=3). (d) Optical density change at 405 nm of hydrolysed product of 4-nitrophenyl N-acetyl-β-D-glucosaminide under pH 10 bathed at 60 °C for 5 h. (ΔOD=OD_time_-OD_0 h_) Data are presented as mean ± standard error of the mean (n=3). (e) GPC profiles of carboxymethyl chitosan before and after treatment with Ce-FMA-FA-20-RT at 37 °C or 60 °C under pH 8.0.

Generally, Ce-FMA-FA-20-RT prefers to cleave polysaccharides with (1) N-acetyl groups; (2) excellent leaving groups (such as 2-nitrophenyl and 4-nitrophenyl). The above findings revolutionize the traditional concept of the acid/base-catalysed cleavage of glycosidic bonds, where glycosidase acts on the carboxyl groups of aspartic acid/glutamate residues. In addition to the abovementioned hard-soft-acid-base theory, we also find the importance of acetyl amino group which forms H-bonding and thereby provides a vicinity for Ce cluster and substrates. Certainly, this new understanding of the MOF-catalysed hydrolysis of glycosidic bonds will expand the product scope beyond that of natural glycosidases and therefore yield novel polysaccharides.

### Application of Ce-FMA-FA-20-RT in hydrolysis of multiple substrates for biofilm degradation

As described above, Ce-FMA-FA-20-RT has been shown to hydrolyse several individual substrates containing phosphoester bonds, amide/peptide bonds, and glycosidic bonds. A more challenging feat is the degradation of mixture of these substrates, such as biofilms. A biofilm is a microbial consortium with self-produced extracellular polymeric substances (EPS). Inside the three dimensional architecture of a biofilm, the EPS forms a scaffold that hosts the bacteria and is responsible for external defence, adhesion to surfaces, connectivity and nutrient trapping. Due to the roles that a biofilm plays as a protector and energy supplier, cells in a biofilm have adopted properties different from those of planktonic bacteria, thus limiting the efficacy of antimicrobials against bacterial infection involving biofilms^48^. One of the reasons why biofilms are so difficult to deal with is their varied compositions: polysaccharides/phosphoethanolamine cellulose^49^, proteins, nucleic acids, and lipids (Figure 6a). This complexity increases the difficulty for a single enzyme to disrupt a biofilm. In this regard, Ce-FMA-FA-20-RT, with multiple hydrolytic activities, may be advantageous, as it has been proven to be effective in cleaving phosphoester bonds, amide/peptide bonds, and glycosidic bonds, which all exist in biofilms in various forms.

To test this hypothesis, we applied two representative biofilms formed by gram-negative bacteria (*E. coli*) and gram-positive bacteria (*S. aureus*). Bacterial cells were grown for 48 h in a 24-well plate stationarily to form biofilms. Then, Ce-FMA-FA-20-RT together with fresh medium was added, followed by another 12 h incubation at 37 °C to allow catalytic hydrolysis of the biofilms. The scanning electronic microscopy (SEM) images in Figure 6b and 6d display the expected thick, bulk-like adhesions between cells when no Ce-FMA-FA-20-RT treatment was applied, demonstrating the successful formation of biofilms. In contrast, the adhesion among cells became tenuous and fibre-like after Ce-FMA-FA-20-RT treatment, and clear gaps were also observed. Crystal violet staining assay was applied to semi-quantitate the amount of biofilm. As shown in Figure 6c and 6e, a significant difference was observed before and after Ce-FMA-FA-20-RT treatment in both gram-negative bacteria (*E. coli*) and gram-positive bacteria (*S. aureus*). Moreover, plates spread with Ce-FMA-FA-20-RT-treated *E. coli* and *S. aureus* indicated negligible adverse effects on bacterial growth, as shown in Figure S19. This finding is also consistent with the intact cell morphology shown in Figure 6b and 6d after treatment with Ce-FMA-FA-20-RT, confirming that the decrease in biofilm formation could only be attributed to hydrolysis rather than to bacterial cell death.

**Figure 6.**
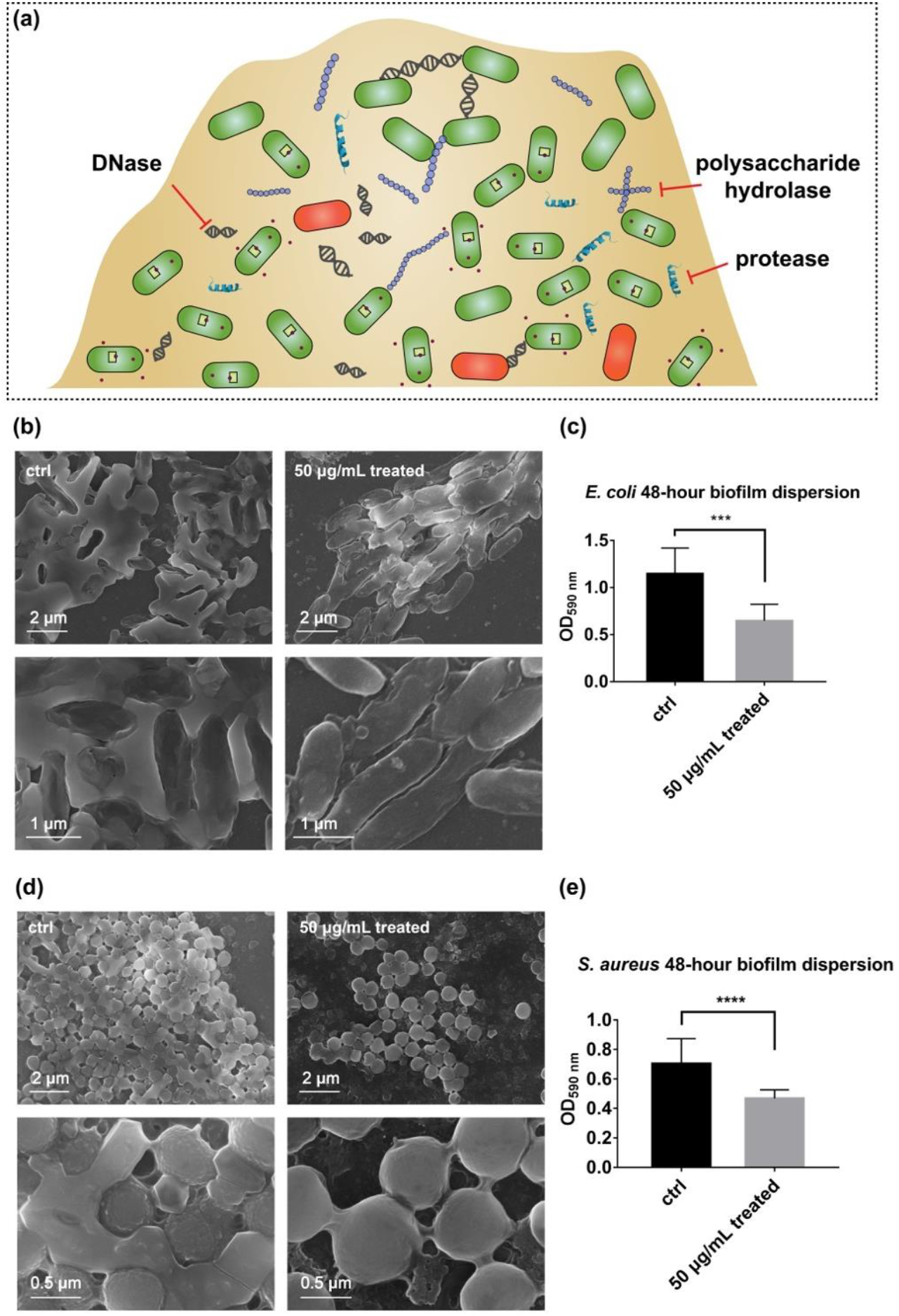
Hydrolytic performance of Ce-FMA-FA-20-RT on biofilms. (a) Schematic of the diverse biofilm components and strategies to combat biofilm formation by using hydrolytic enzymes. (b) SEM images of biofilms formed by *E. coli* without/with Ce-FMA-FA-20-RT treatment (upper: low magnification, lower: high magnification). (c) Crystal violet staining of biofilms formed by *E. coli* without/with Ce-FMA-FA-20-RT treatment (significance level: ***, P<0.001). Data are presented as the mean ± standard error after removing the maximum and minimum, n=12. (d) SEM images of biofilms formed by *S. aureus* without/with Ce-FMA-FA-20-RT treatment (upper: low magnification, lower: high magnification). (e) Crystal violet staining of biofilms formed by *S. aureus* without/with Ce-FMA-FA-20-RT treatment (significance level: ****, P<0.0001). Data are presented as the mean ± standard error after removing the maximum and minimum, n=16.

## Conclusions

In conclusion, we introduced a data-informed approach to discover a high performance hydrolytic nanozyme by systematically analysing 105 published papers to supplement the ill-developed field of research on hydrolytic nanozymes. Analysis of these data indicates that MOF is a good scaffold to embed hydrolytic active sites for their tuneable metal clusters and ligands. Further structured data suggest that Lewis acidity of metal clusters in MOFs and the volume density of active sites which is adjusted by ligand are two critical elements. Consequently, we screened Ce^4+^ as a promising metal ion component and applied a short ligand, FMA, to construct a UiO-66–like MOF.

Encouragingly, this rationally designed Ce-FMA was highly efficient in hydrolysing a broad scope of substrates and exhibited huge promise to be further developed. First, nonactivated Ce-FMA exhibited excellent phosphatase-like activity (half-life of less than 2 min) even without the use of co-catalysts. In addition to widely researched organophosphorus antidote, it is also promising to apply ALP-like nanozyme in bone regeneration as ALP can transform pyrophosphate to free phosphate and thus accelerate mineralization in hard tissue formation. Second, Ce-FMA showed high activity towards BSA hydrolysis, with 12.7 times better efficiency than the multi-functional Zr-based MOF-808. The new cleavage sites (such as D sites) brought by nanozymes expand the diversity of peptides and offer additional probability to obtain active peptides. What is more, the high environmental tolerance and stability of nanozymes provide feasibility to degrade plastics such as nylon (a polymer which is linked by amide bonds) and thereby to upcycle the unsustainable the plastic. Third, we evaluated the ability of this Ce-FMA MOF to cleave glycosidic bonds, advancing the understanding of traditional acid/base-accelerated reactions and enriching the product scope beyond those of natural glycosidases. Last, we applied Ce-FMA to a mixture of biomacromolecules, *i.e.*, biofilms, from both gram-negative bacteria (*E. coli*) and gram-positive bacteria (*S. aureus*), which is hopefully applied in biomedicine and marine. Additionally, this data-informed approach can be applied to deduce abiotic active sites rather than being limited to the direct modification of natural active sites.

We acknowledge that more chemical information about hydrolytic nanozymes should be analysed, such as the topology of the coordination network, the coordination number of metal clusters and the types of functional groups. However, due to the limited amount of available data, the current study was not suitable for machine learning or chemoinformatic models with multiple variables and was unable to discover esterase mimics. Nevertheless, given the rapid development of this field, we anticipate that machine learning as well as other artificial intelligence techniques will be further leveraged for the design and discovery of new nanozymes, such as powerful esterase mimics for degradation of PET (polyethylene terephthalate) in the near future.

## Acknowledgements

This work was supported by National Natural Science Foundation of China (21722503 and 21874067), the National Key R&D Program of China (2019YFA0709200), PAPD Program, Fundamental Research Funds for the Central Universities (14380145), and Interdisciplinary Project Funded by Graduate School of Nanjing University (2017CL12).

## References

1 Mirts, E. N., Petrik, I. D., Hosseinzadeh, P., Nilges, M. J. & Lu, Y. A designed heme-[4Fe-4S] metalloenzyme catalyzes sulfite reduction like the native enzyme. Science 361, 1098–1101, (2018).

2 Grommet, A. B., Feller, M. & Klajn, R. Chemical reactivity under nanoconfinement. Nat. Nanotechnol. 15, 256–271, (2020).

3 Wu, J. et al. Nanomaterials with enzyme-like characteristics (nanozymes): Next-generation artificial enzymes (II). Chem. Soc. Rev. 48, 1004–1076, (2019).

4 Gao, L. et al. Intrinsic peroxidase-like activity of ferromagnetic nanoparticles. Nat. Nanotechnol. 2, 577–583, (2007).

5 Manea, F., Houillon, F. B., Pasquato, L. & Scrimin, P. Nanozymes: Gold-nanoparticle-based transphosphorylation catalysts. Angew. Chem. Int. Ed. 43, 6165–6169, (2004).

6 Huang, Y., Ren, J. & Qu, X. Nanozymes: Classification, catalytic mechanisms, activity regulation, and applications. Chem. Rev. 119, 4357–4412, (2019).

7 Liang, M. & Yan, X. Nanozymes: From new concepts, mechanisms, and standards to applications. Acc. Chem. Res. 52, 2190–2200, (2019).

8 Wei, H. & Wang, E. Fe_3_O_4_ magnetic nanoparticles as peroxidase mimetics and their applications in H_2_O_2_ and glucose detection. Anal. Chem. 80, 2250–2254, (2008).

9 Zhang, Z., Zhang, X., Liu, B. & Liu, J. Molecular imprinting on inorganic nanozymes for hundred-fold enzyme specificity. J. Am. Chem. Soc. 139, 5412–5419, (2017).

10 Gupta, A., Das, R., Yesilbag Tonga, G., Mizuhara, T. & Rotello, V. M. Charge-switchable nanozymes for bioorthogonal imaging of biofilm-associated infections. ACS Nano 12, 89–94, (2018).

11 Kim, J. et al. Continuous O_2_-evolving MnFe_2_O_4_ nanoparticle-anchored mesoporous silica nanoparticles for efficient photodynamic therapy in hypoxic cancer. J. Am. Chem. Soc. 139, 10992–10995, (2017).

12 Mandoli, C. et al. Stem cell aligned growth induced by CeO_2_ nanoparticles in PLGA scaffolds with improved bioactivity for regenerative medicine. Adv. Funct. Mater. 20, 1617–1624, (2010).

13 Singh, B. K. Organophosphorus-degrading bacteria: Ecology and industrial applications. Nat. Rev. Microbiol. 7, 156–164, (2009).

14 Xia, M. et al. Assembly of the active center of organophosphorus hydrolase in metal–organic frameworks via rational combination of functional ligands. Chem. Commun. 53, 11302–11305, (2017).

15 Jin, C., Zhang, S., Zhang, Z. & Chen, Y. Mimic carbonic anhydrase using metal–organic frameworks for CO2 capture and conversion. Inorg. Chem. 57, 2169–2174, (2018).

16 Pieters, G., Pezzato, C. & Prins, L. J. Controlling supramolecular complex formation on the surface of a monolayer-protected gold nanoparticle in water. Langmuir 29, 7180–7185, (2013).

17 Chen, J. et al. Bio-inspired nanozyme: A hydratase mimic in a zeolitic imidazolate framework. Nanoscale 11, 5960–5966, (2019).

18 Maiti, S., Fortunati, I., Ferrante, C., Scrimin, P. & Prins, L. J. Dissipative self-assembly of vesicular nanoreactors. Nat. Chem. 8, 725–731, (2016).

19 Sun, M. et al. Site-selective photoinduced cleavage and profiling of DNA by chiral semiconductor nanoparticles. Nat. Chem. 10, 821–830, (2018).

20 Vaissier Welborn, V. & Head-Gordon, T. Computational design of synthetic enzymes. Chem. Rev. 119, 6613–6630, (2019).

21 Key, H. M., Dydio, P., Clark, D. S. & Hartwig, J. F. Abiological catalysis by artificial haem proteins containing noble metals in place of iron. Nature 534, 534–537, (2016).

22 Zhou, Z. & Roelfes, G. Synergistic catalysis in an artificial enzyme by simultaneous action of two abiological catalytic sites. Nat. Catal., 289–294, (2020).

23 Wilhelm, S. et al. Analysis of nanoparticle delivery to tumours. Nat. Rev. Mater. 1, 16014, (2016).

24 Yaghi, O. M. et al. Reticular synthesis and the design of new materials. Nature 423, 705–714, (2003).

25 Zhao, M. et al. Metal–organic frameworks as selectivity regulators for hydrogenation reactions. Nature 539, 76–80, (2016).

26 Horike, S., Shimomura, S. & Kitagawa, S. Soft porous crystals. Nat. Chem. 1, 695, (2009).

27 Mondloch, J. E. et al. Destruction of chemical warfare agents using metal–organic frameworks. Nat. Mater. 14, 512–516, (2015).

28 Son, F. A. et al. Uncovering the role of metal–organic framework topology on the capture and reactivity of chemical warfare agents. Chem. Mater. 32, 4609–4617, (2020).

29 Park, H. J. et al. Synthesis of a Zr-based metal–organic framework with spirobifluorenetetrabenzoic acid for the effective removal of nerve agent simulants. Inorg. Chem. 56, 12098–12101, (2017).

30 Ly, H. G. T. et al. Superactivity of MOF-808 toward peptide bond hydrolysis. J. Am. Chem. Soc. 140, 6325–6335, (2018).

31 Feng, D. et al. Zirconium-metalloporphyrin PCN-222: Mesoporous metal–organic frameworks with ultrahigh stability as biomimetic catalysts. Angew. Chem. Int. Ed. 51, 10197–10197, (2012).

32 Feng, D. et al. A highly stable zeotype mesoporous zirconium metal–organic framework with ultralarge pores. Angew. Chem. Int. Ed. 54, 149–154, (2015).

33 Liu, W.-G. & Truhlar, D. G. Computational linker design for highly crystalline metal–organic framework NU-1000. Chem. Mater. 29, 8073–8081, (2017).

34 Peterson, G. W. et al. Tailoring the pore size and functionality of UiO-type metal–organic frameworks for optimal nerve agent destruction. Inorg. Chem. 54, 9684–9686, (2015).

35 Hu, Z. et al. Modulator effects on the water-based synthesis of Zr/Hf metal–organic frameworks: Quantitative relationship studies between modulator, synthetic condition, and performance. Cryst. Growth Des. 16, 2295–2301, (2016).

36 Liu, L. et al. Imaging defects and their evolution in a metal–organic framework at sub-unit-cell resolution. Nat. Chem. 11, 622–628, (2019).

37 Wang, S., McGuirk, C. M., d'Aquino, A., Mason, J. A. & Mirkin, C. A. Metal–organic framework nanoparticles. Adv. Mater., e1800202, (2018).

38 Islamoglu, T. et al. Cerium(IV) vs zirconium(IV) based metal–organic frameworks for detoxification of a nerve agent. Chem. Mater. 29, 2672–2675, (2017).

39 Lammert, M. et al. Cerium-based metal organic frameworks with UiO-66 architecture: Synthesis, properties and redox catalytic activity. Chem. Commun. 51, 12578–12581, (2015).

40 Plonka, A. M. et al. In situ probes of capture and decomposition of chemical warfare agent simulants by Zr-based metal organic frameworks. J. Am. Chem. Soc. 139, 599–602, (2017).

41 Katz, M. J. et al. One step backward is two steps forward: Enhancing the hydrolysis rate of UiO-66 by decreasing [OH^−^]. ACS Catal. 5, 4637–4642, (2015).

42 Radzicka, A. & Wolfenden, R. Rates of uncatalyzed peptide bond hydrolysis in neutral solution and the transition state affinities of proteases. J. Am. Chem. Soc. 118, 6105–6109, (1996).

43 de Koning, M. C., van Grol, M. & Breijaert, T. Degradation of Paraoxon and the chemical warfare agents VX, Tabun, and Soman by the metal–organic frameworks UiO-66-NH_2_, MOF-808, NU-1000, and PCN-777. Inorg. Chem. 56, 11804–11809, (2017).

44 Li, A. et al. Chemical cleavage at aspartyl residues for protein identification. Anal. Chem. 73, 5395–5402, (2001).

45 Swaney, D. L., Wenger, C. D. & Coon, J. J. Value of using multiple proteases for large-scale mass spectrometry-based proteomics. J. Proteome Res. 9, 1323–1329, (2010).

46 Garrido-González, J. J. et al. An enzyme model which mimics chymotrypsin and N-terminal hydrolases. ACS Catal. 10, 11162–11170, (2020).

47 Walther, R. et al. Identification and directed development of non-organic catalysts with apparent pan-enzymatic mimicry into nanozymes for efficient prodrug conversion. Angew. Chem. Int. Ed. 58, 278–282, (2019).

48 Flemming, H. C. et al. Biofilms: An emergent form of bacterial life. Nat. Rev. Microbiol. 14, 563–575, (2016).

49 Thongsomboon W. et al. Phosphoethanolamine cellulose: A naturally produced chemically modified cellulose. Science 359, 334–338, (2018).

